# Impulsivity and Thought Suppression in Behavioral Addiction: Associated Neural Connectivity and Neural Networks

**DOI:** 10.1101/2020.10.14.340273

**Authors:** Li Wan, Rujing Zha, Jiecheng Ren, Ying Li, Qian Zhao, Huilin Zuo, Xiaochu Zhang

## Abstract

Impulsivity and thought suppression are two psychological traits that have great variation in healthy population. In extreme cases, both are closely related to mental illness and play an important role in behavioral addiction. We have known the role of the top-down mechanism in impulsivity and thought suppression, but we do not know how the related neural nuclei are functionally connected and interact with each other. In the study, we selected excessive internet users (EIU) as our target population and investigated the relationship between thought suppression and impulsivity in the following aspects: their correlations to psychological symptoms; the associated neural networks; and the associated brain morphometric changes. We acquired data from 131 excessive internet users, with their psychological, resting-state fMRI and T1-MRI data collected. With the whole brain analysis, graph theory analysis, replication with additional brain atlas, replication with additional MRI data, and analysis of brain structure, we found that: (i) implusivity and thought suppression shared common neural connections in the top-down mechanism; (ii) thought suppression was associated with the neural network that connected to the occipital lobe in the resting-state brain but not the morphometric change of the occipital lobe. The study confirmed the overlap between impulsivity and thought suppression in terms of neural connectivity and suggested the role of thought suppression and the occipital network in behavioral addiction. Studying thought suppression provided a new insight into behavioral addiction research. The neural network study helped further understanding of behavioral addiction in terms of information interaction in the brain.

## INTRODUCTION

### 1. Impulsivity and Thought Suppression in Behavioral Addiction

Since the widespread use of the internet, internet addiction or excessive internet use has become health issues that need attention, especially among teenagers and young adults. For internet addiction, the key problem is in the combination of decision-making style (choose immediate rather than long-term reward) and motivation seeking (pleasure pursuit and pressure reduction), and ultimately match the patient’s executive and control functions ^1^. This view was consistent with the main theories in substance addiction ^2^. Brand et al ^3^ proposed another theory, internet addiction was due to an individual’s susceptibility to executive and inhibitory functions, a tendency to cognitive and emotional biases, and obvious biases in decision-making (tend to be immediate satisfaction), so addictive behavior can be maintained or even strengthened. Impaired control of onset, frequency, duration, intensity, termination and context of internet use have been consistently found in excessive internet users ^4^.

Impulsivity and thought suppression are two psychological traits that have great variation in healthy people ^5, 6^. In extreme cases, both are closely related to mental illness. Especially, both play an important role in addiction. Impulsivity was defined as “action that is inappropriate, expresses prematurely, is too risky or is inappropriate to the situation, and often leads to undesirable consequences” ^7^. Impulsivity is a symptom dimension in psychiatry ^8^ and manifests as a series of mental disorders, including obsessive-compulsive disorder (OCD) ^9^, attention deficit/hyperactivity disorder (ADHD) ^10^, substance abuse ^11^, addictive disorders ^12^, aggressive behavior ^13^, and manic and anti-social behavior ^14^.

Thought suppression was the suppression of intrusive thinking, which was involuntary memories, images, ruminations or hallucinations that entered mind without an expectation ^5^. Intrusive thinking was usually triggered by specific cues and then led to symptoms or the recurrence of undesirable behaviors ^15^. However, simply suppressing intrusive thoughts actually lead to higher level and higher frequency of intrusive thinking, compared to monitoring thoughts without suppressing them ^15–17^. Maladaptive intrusive thinking also caused psychiatric disorders, such as anxiety ^18^, depression ^19^, obsessive-compulsive disorder (OCD) ^20^, schizophrenia ^21^, posttraumatic stress disorder (PTSD) ^22^, and substance abuse ^17^. The levels of thought suppression were positively correlated with anxiety traits ^23, 24^, generalized anxiety disorders ^25^, OCD ^26^, obsessive thinking ^23^, depressive symptoms ^27^, substance abuse ^16^, and addiction ^17^.

### 2. Neuroimaging Findings on Impulsivity and Thought Suppression

The neuroimaging studies have largely revealed the neural nuclei and neural mechanisms related to impulsivity and thought suppression.

The main components of the impulsive neural matrix have been found to include the nucleus accumbens (NAcc), inferior and anterior prefrontal cortex (PFC), anterior cingulate cortex (ACC), and hippocampus ^28,29^. Impulsivity is mediated by neural circuits related to motivation and decision-making processes, including the basal ganglia, its border cortical inputs, and the top-down control of the cortical prefrontal circuits ^30–32^. More specifically, the reward reduction circuit included the ventromedial PFC, the inferior cingulate cortex, NAcc, and ventral striatum. The motor inhibitory circuit includes the ventrolateral PFC, ACC, complementary movement cortex and their connection with the caudate and putamen ^33^. A recent study reported that the reduction of the myelin-sensitive magnetization transfer trajectory in the frontal foramen region was associated with the expression of impulsivity, the reduction was most pronounced in the lateral and medial PFC areas ^34^.

Damage to the cortico-striatal nervous system largely contributed to maladaptive invasive thinking. Imaging studies have found that addiction-related cues increased the activation in the ACC, amygdala, ventral striatum, NAcc in addictive disorders, such as drug use ^35^, gambling ^36^, and overeating ^37^. On the contrary, increased activation in ACC and insular during the suppression of thoughts were found in comparison with free-thought conditions ^38^. The activities of the medial PFC, ACC and left lateral PFC could be triggered by suppression of thoughts ^39^. The right dorsolateral PFC and ACC were responsible for sustained and transient suppression ^40^. In depressed patients, reduced activation in the right middle frontal gyrus and increased activation in the amygdala and hippocampus were associated with memory suppression involving negatively valenced stimuli ^41^. Reduced activation in the inferior frontal gyrus and supramarginal gyri were related to though suppression ^42^.

Neuroimaging studies have found that impulsivity and thought suppression were both regulated by the top-down mechanism, including the PFC, subcortical and limbic structures as well as the PFC-striatal projection. Under the pathological condition, the reduced PFC activities and enhanced subcortical activities lead to the occurrence of intrusive thinking and/or impulsivity.

### 3. The Purpose of the Study

Brain regions (or neural nuclei) did not exist in isolation, there was always information interaction between brain regions and the brain exist and operated in the forms of networks ^43^. The connections between brain regions in the resting state were relatively stable, forming the basis of cognition and behavior ^44^, and certain neural networks showed unique changes in mental illness ^45,46^. We have known the role of the top-down mechanism in impulsivity and thought suppression, but we do not know how the related neural nuclei are functionally connected and interact with each other. The neural network study will provide new insights into the understanding of psychiatric disorders in terms of information interaction in the brain.

In the study, we selected excessive internet users (EIU) as our target population, we investigated the relationship between thought suppression and impulsivity in EIU in the following aspects: (i) their correlations to psychological measures including psychological symptoms; (ii) the associated neural networks; and (iii) the associated brain morphometric changes. In the study, we integrated multiple levels of information from gene to brain imaging to psychological measures. The work was registered in the Open Science Foundation (https://osf.io).

## METHODS AND MATERIALS

The experimental approach was summarized in Figures 1 and 2. For more details, also see supplementary experimental procedure.

**Figure 1:**
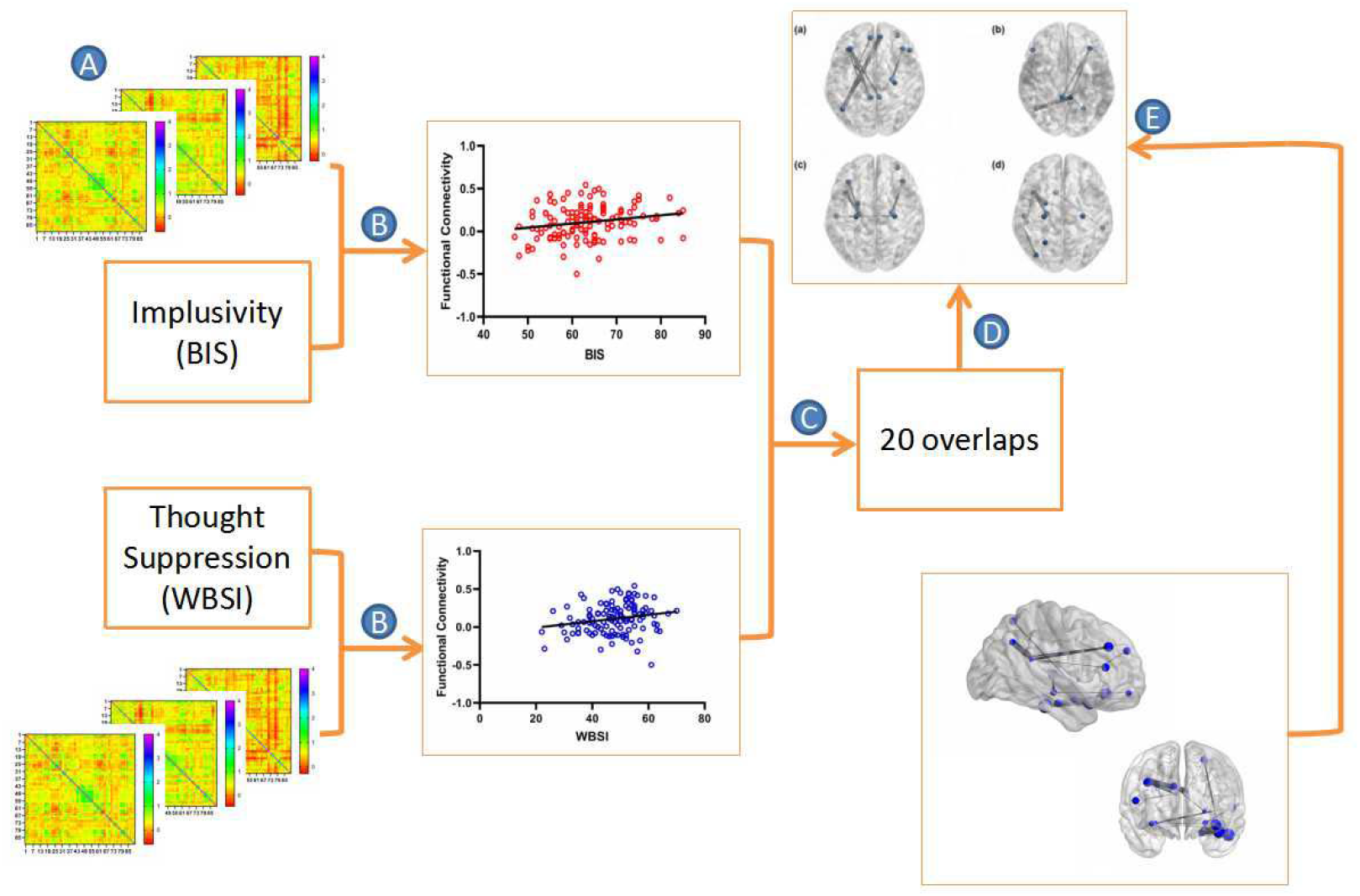
Summary of methods. A. calculated functional connectivity between any two nodes (90×90 on the AAL template). B. Pearson’s correlation between the connectivity values and BIS or WBSI scores. C. connections significantly related to both BIS and WBSI scores were the overlap. D. grouped the 20 connections into 4 categories based on the anatomic locations. E. using the Brainnetome template (246×246 nodes) to verify the overlapped connections.

**Figure 2:**
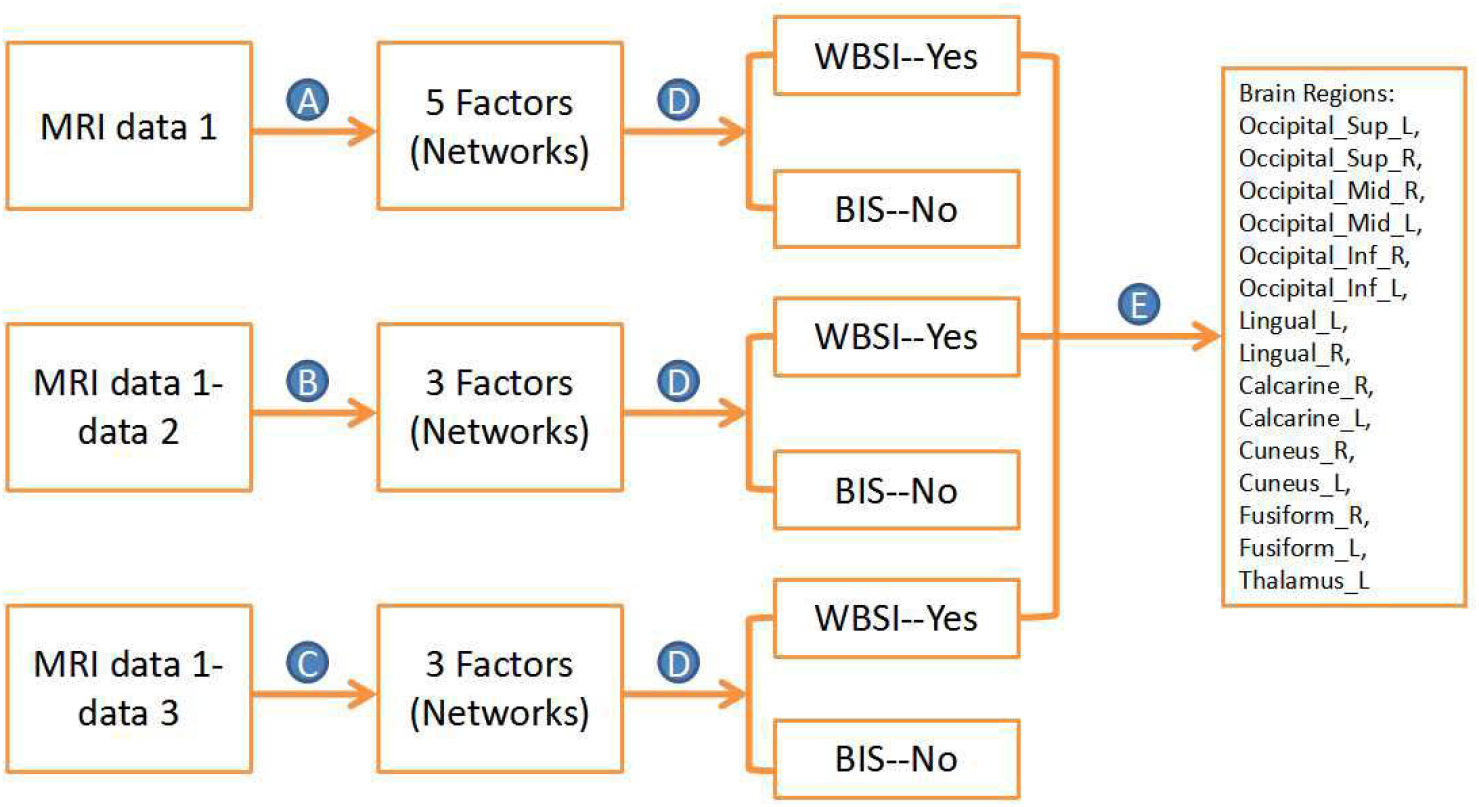
Summary of methods. A. the factor analysis concentrated 90 nodes to 5 factors (groups of nodes or networks). B. the factor analysis concentrated 38 nodes to 3 factors. C. the factor analysis concentrated 48 nodes to 3 factors. D. the factor score for each factor was calculated and correlated to BIS/WBSI total score seperately. E. correlations with factor scores (or network scores) were found for WBSI but not for BIS. Brain regions that were involved in the results of all three analyses were extracted. MRI data 1: EIU data; data 2: HCP data; data 3: FCP data.

### 1. Participants

Data were acquired from 131 internet users. Participants who used internet for at least 12 months and continued internet use daily in the past 3 months, beyond the purpose of work or study, were recruited. They had no history of current neurological or psychiatric disorders, including substance use and no medicine use or substance abuse in the past 3 months.

### 2. Psychological Measures

Psychological traits were measured with Excessive Internet Use (EIU) ^47^, Symptom Check list 90-Revised (SCL-90R) ^48^, Barratt Impulsiveness Scale (BIS) ^49^, White Bear Suppression Inventory (WBSI) ^23^, World Health Organization Quality of Life Scale-Brief Form (WHO-QoL) ^50^, and Tridimensional Personality Questionnaire (TPQ) ^51^.

### 3. Magnetic Resonance Imaging

MRI data were acquired using two identical 3-T Siemens Magnetom Trio scanners (Siemens, Erlangen, Germany). Functional images were acquired with a T2-weighted gradient echo-planar imaging sequence (TR=2s, TE=30ms, FOV=240mm, matrix, 64×64) with 33 axial slices (3.7mm thickness, no gap), covering the whole brain. High resolution T1-weighted spin-echo images were also collected for anatomical overlay. Resting-state fMRI data were acquired with a 8-min run (240 epochs).

Additional data were acquired from two sources: the Human Connectome Project (HCP; http://humanconnectome.org), and the 1000 Functional Connectomes Project (FCP, http://fcon_1000.projects.nitrc.org).

### 4. Identifying Associated Neural Connectivity

The imaging data were processed with the Analysis of Functional Neuroimages (AFNI). Using the graphical theory analysis ^52^ and the automated anatomical labeling atlas (AAL; ^53^, we divided the whole brain into 90 brain regions (or nodes) to calculate the functional connectivity between any two nodes. For repetition and verification, we have used another atlas and obtained 246×246 functional connection matrixbased on the partitions of the Brainnetome atlas ^54^, the correlation coefficient of any two nodes was calculated with 237 points on the time course, and the corresponding z value after the fisher-z transformation of the correlation coefficient was calculated. Pearson’s correlations were conducted between these values and BIS or WBSI scores separately.

### 5. Identifying Associated Neural Networks

The strength of each node (AAL 90 nodes) was then calculated by averaging the connectivity to other 89 nodes. To generate the networks, we conducted an exploratory factor analysis with the method used in a previous study ^55^. The factor analysis concentrated 90 nodes to a smaller number of factors (or groups of nodes) based on their relative strength. The factor scores were then correlated with BIS and WBSI scores separately with Pearson’s correlation (p < 0.05).

For repetition and verification, we used additional MRI data.We first compared the strength values of 90 nodes among the three samples and extracted the nodes that were significant different. We then conducted factor analyses with the extracted nodes. The factor scores obtained were then correlated with the scores of BIS and WBSI respectively.

### 6. Identifying Associated Brain Morphometric Changes

Cortical surface reconstruction was performed with standard procedures provided by the software (http://surfer.nmr.mgh.harvard.edu/fswiki). We obtained an array of anatomical measures, including: cortical thickness, surface area, and curvature at each region on the cortex, within the Desikan-Killiany atlas ^56^. Pearson’s correlations were conducted between these indexes and BIS/WBSI scores separately.

## RESULTS

One hundred and thirty-one participants took part in the study, mean age = 19.74, SD = 2.83, all males.

### 1. Psychological Measures

The mean and standard deviation of each item in the questionnaires were presented in Table S1 and S2. Pearson’s correlation between BIS and WBSI total scores was 0.38, p < 0.001, indicating an overlap between impulsivity and thought suppression. In addition, both impulsivity and thought suppression were associated with nine psychological symptoms, including somatization. obsessive-compulsive symptoms, interpersonal sensitivity, depression, anxiety, hostility, terror, paranoid, and psychosis. The BIS and WBSI total scores were positively correlated with the scores of the nine items in SCL-90R (Table 1). The BIS total score was positively correlated with the internet use score of the past one year, and the WBSI score was positively correlated with the internet use score of the most severe week in the past one year (Table 1).

**Table 1:** The Pearson’s correlations between BIS/WBSI total score and other psychological measures (n = 131).

|  | Items | BIS total score | WBSI total score |  |
| --- | --- | --- | --- | --- |
| **Quality of life** | Physiology | -.30** | -.01 | ** p < 0.01<br>* p < 0.05 |
|  | Psychology | -.38** | -.23* |  |
|  | Environment | -.38** | -.41** |  |
| **Personality** | Curiosity | .47** | .04 |  |
|  | Avoidance of harm | .29** | .30** |  |
|  | Reward dependence | -.15 | .01 |  |
| **Excessive internet use** | In the most severe week | .14 | .19* |  |
|  | In the past year | .22* | -.11 |  |
| **Symptom checklist** | Somatization | .33** | .27** |  |
|  | Obsessive-compulsive | .53** | .43** |  |
|  | Interpersonal sensitivity | .46** | .46** |  |
|  | Depression | .48** | .41** |  |
|  | Anxiety | .41** | .38** |  |
|  | Hostility | .34** | .33** |  |
|  | Terror | .43** | .39** |  |
|  | Paranoid | .44** | .45** |  |
|  | Psychosis | .39** | .48** |  |

### 2. Identifying the Associated Neural Connectivity

Among 90 × 90 connections, 189 connections significantly correlated with BIS total score and 280 connections significantly correlated with WBSI total score. Among these correlations, 20 were correlated with both BIS and WBSI scores in same directions (Table 2 and Table S3). The higher level of thought suppression and impulsivity were associated with stronger connectivity value in the 20 connections. The 20 connections were divided into 4 groups based on the anatomic locations: the frontal lobe related connectivity (n = 9), the temporal lobe related connectivity (n = 8), the hippocampus related connectivity (n = 10), and the cingulum related connectivity (n = 5) (Table 2 and Figure 3).

**Table 2.**
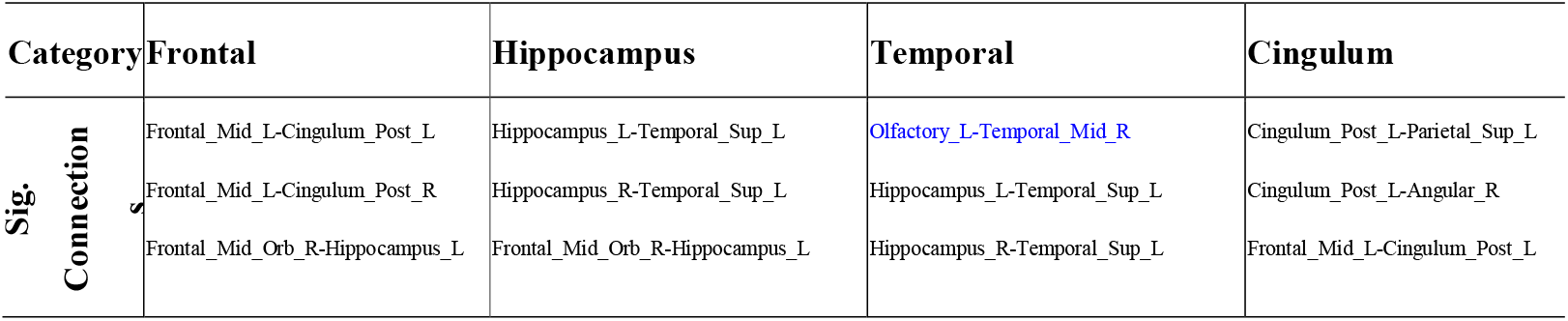
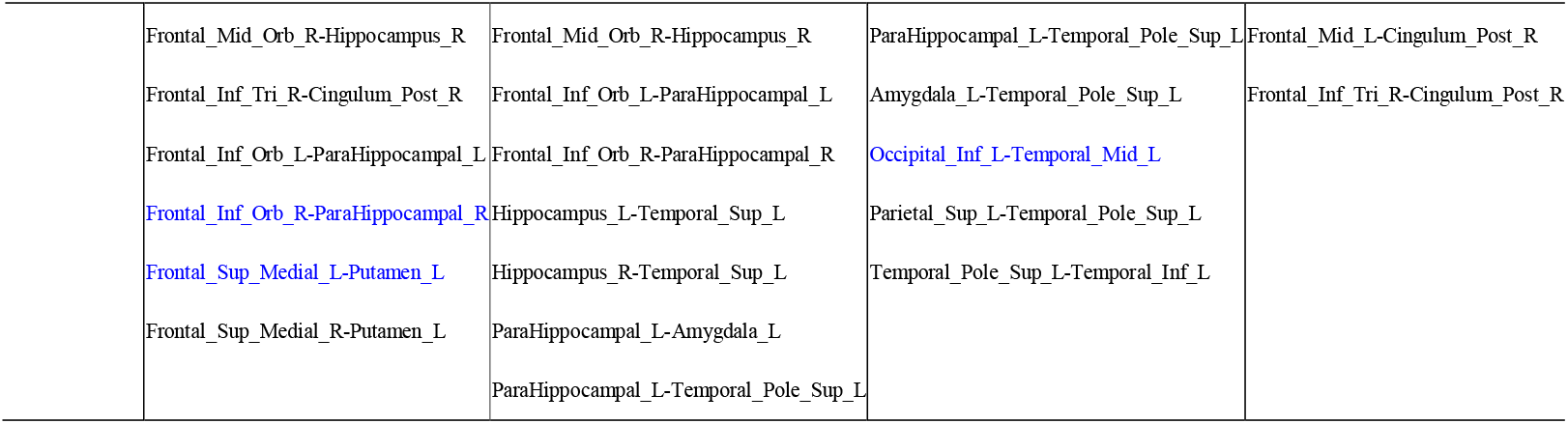
The neural connections associated to thought suppression and impulsivity, identified by the AAL template (in black were confirmed by the Brainnetome template, in blue were not confirmed).

| Category | Frontal | Hippocampus | Temporal | Cingulum |
| --- | --- | --- | --- | --- |
| **Sig. Connection** | Frontal_Mid_L-Cingulum_Post_L | Hippocampus_L-Temporal_Sup_L | Olfactory_L-Temporal_Mid_R | Cingulum_Post_L-Parietal_Sup_L |
|  | Frontal_Mid_L-Cingulum_Post_R | Hippocampus_R-Temporal_Sup_L | Hippocampus_L-Temporal_Sup_L | Cingulum_Post_L-Angular_R |
|  | Frontal_Mid_Orb_R-Hippocampus_L | Frontal_Mid_Orb_R-Hippocampus_L | Hippocampus_R-Temporal_Sup_L | Frontal_Mid_L-Cingulum_Post_L |

|  |  |  |  |
| --- | --- | --- | --- |
| Frontal_Mid_Orb_R-Hippocampus_R | Frontal_Mid_Orb_R-Hippocampus_R | ParaHippocampal_L-Temporal_Pole_Sup_L | Frontal_Mid_L-Cingulum_Post_R |
| Frontal_Inf_Tri_R-Cingulum_Post_R | Frontal_Inf_Orb_L-ParaHippocampal_L | Amygdala_L-Temporal_Pole_Sup_L | Frontal_Inf_Tri_R-Cingulum_Post_R |
| Frontal_Inf_Orb_L-ParaHippocampal_L | Frontal_Inf_Orb_R-ParaHippocampal_R | Occipital_Inf_L-Temporal_Mid_L |  |
| Frontal_Inf_Orb_R-ParaHippocampal_R | Hippocampus_L-Temporal_Sup_L | Parietal_Sup_L-Temporal_Pole_Sup_L |  |
| Frontal_Sup_Medial_L-Putamen_L | Hippocampus_R-Temporal_Sup_L | Temporal_Pole_Sup_L-Temporal_Inf_L |  |
| Frontal_Sup_Medial_R-Putamen_L | ParaHippocampal_L-Amygdala_L |  |  |
|  | ParaHippocampal_L-Temporal_Pole_Sup_L |  |  |

**Figure 3:**
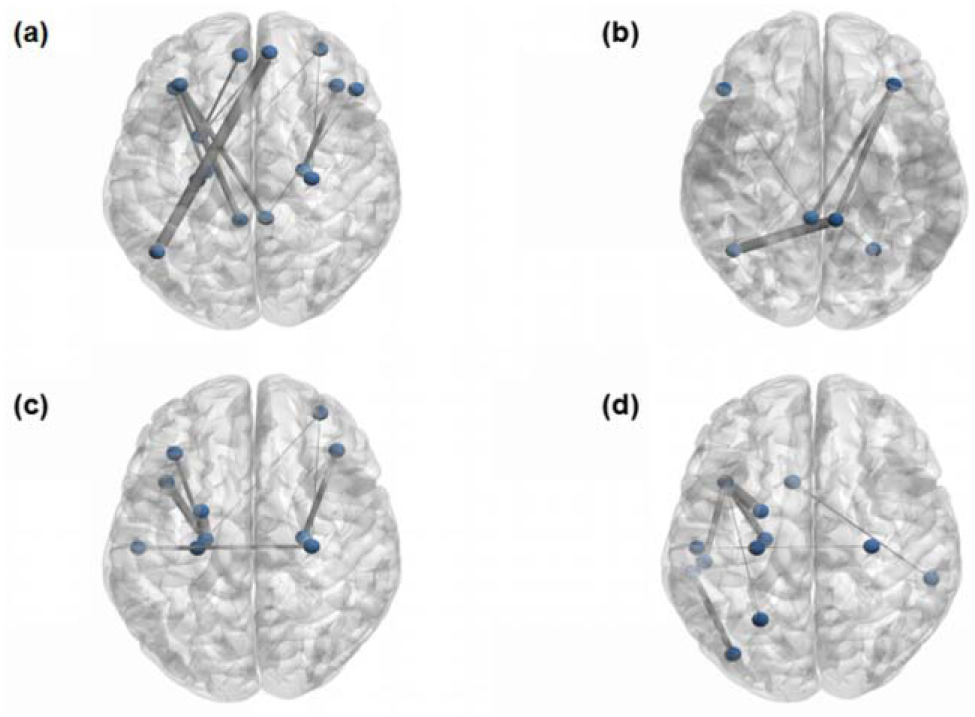
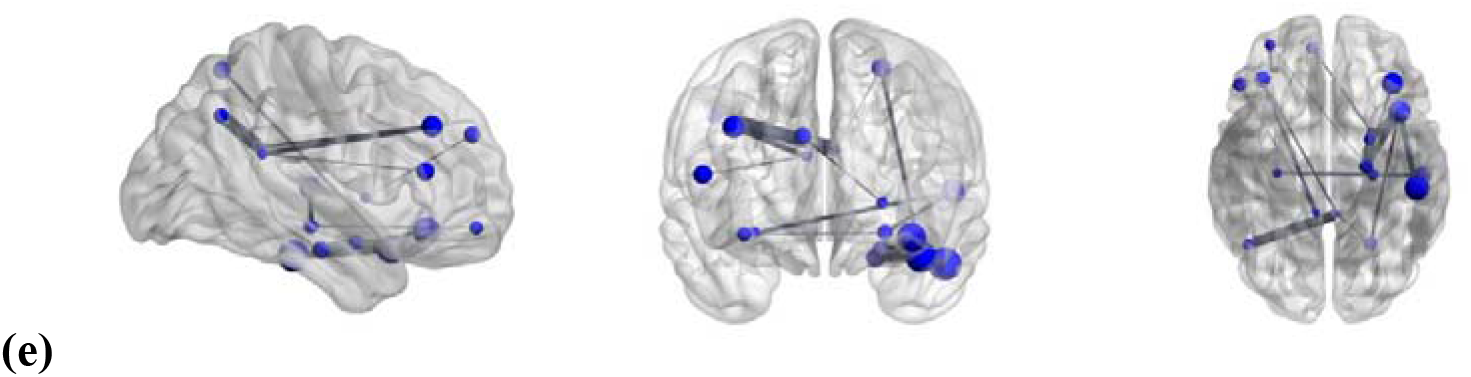
The 20 connections were divided into four categories: a) the frontal; b) the cingulum; c) the hippocampus; d) the temporal. They were correlated to both thought suppression and impulsivity, ps < 0.05. e) The 16 connections confirmed by two brain templates: AAL and Brainnetome.

For repetition and verification, we obtained 246×246 functional connection matrix of each subject according to the partitions of the Brainnetome atlas. Among 20 connections identified above, 16 were confirmed and repeated to be significantly correlated to BIS and WBSI scores (Table 2 and Figure 3).

### 3. Identifying the Associated Neural Networks

#### 3.1 Node Strength and Factor Analysis

The factor analysis with 90 node strengthness revealed spatially separated but functionally related brain regions grouped in the same network. The factors had item loadings of 0.5 or greater, all nodes had a commonality over 0.45 following extraction. The Kaiser-Meyer-Olkin (KMO) measure was 0.898, indicating underlying correlation between variables (nodes). Bartlett’s test of sphericity was significant (χ^2^(4005) =16347, P < 0.001), indicating an underlying correlation structure.

The factor analysis identified 5 major factors (groups of nodes) for resting-state brain activities (Table 2): factor 1. the frontal-cingulum I; factor 2. the occipital; factor 3. the temporal and parietal; factor 4. the frontal-cingulum II; and factor 5. the thalamus. According to the factor loading matrix of each network, the factor scores were calculated with a regression method that weighted each node score.

The factor scores were then correlated to WBSI and BIS scores separately. Each correlation was controlled for age, education, handedness, and scan site.The WBSI total score was positively correlated with factor 4 score (Pearson’s r = 0.27, p < 0.05), and negatively correlated with factor 2 score (Pearson’s r = −0.22, p < 0.05). Greater thought suppression was associated with a lower activity (or connectivity) in the occipital network and a higher activity in the frontal-cingulum II network (Figure 4). There was no significant correlation between BIS scores with the factor scores, ps > 0.05.

**Figure 4:**
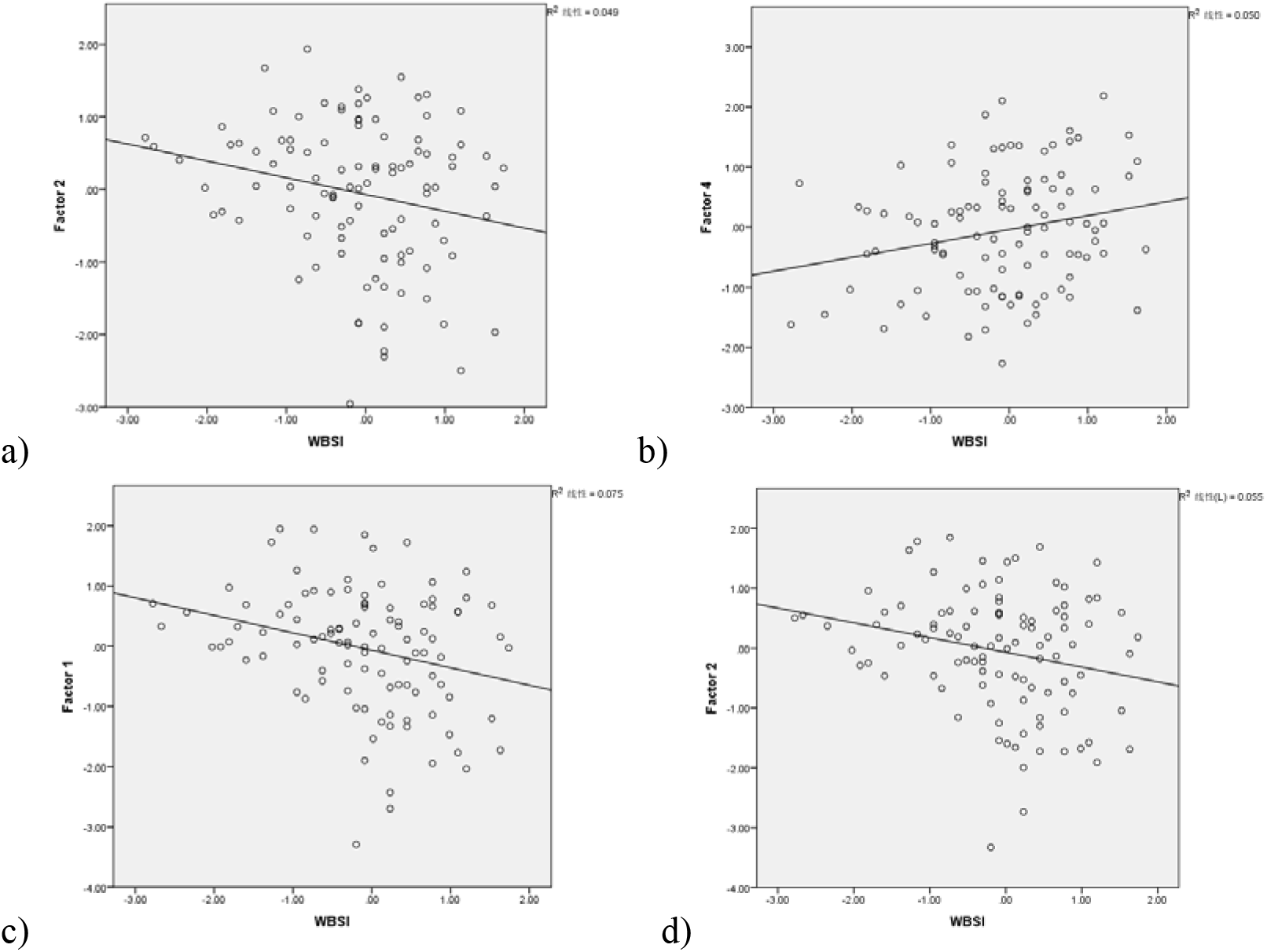
Correlation between the factor score of the network and WBSI score: a) the occipital network (from EIU data only); b) the frontal-cingulum II network (from EIU data only); c) the occipital network (from EIU & HCP data); d) the occipital network (from EIU & FCP data), ps < 0.05.

#### 3.2 Duplication and Confirmation

To duplicate the findings, we used additional MRI data and calculated the strength values for each participant. With independent sample t-tests (p < 0.05), EIU participants had greater node strength values than HCP participants on 38 nodes; EIU participants had greater node strength values than FCP participants on 48 nodes.

The factor analysis with the 38 nodes identified 3 major neural networks: factor 1. the occipital; factor 2. the frontal; and factor 3. the temporal. The WBSI total score was negatively correlated with the factor score of the occipital network (Pearson’s r = −0.27, p < 0.05). There was no significant correlation between BIS scores with the networks, ps > 0.05 (Table 3).

**Table 3:**
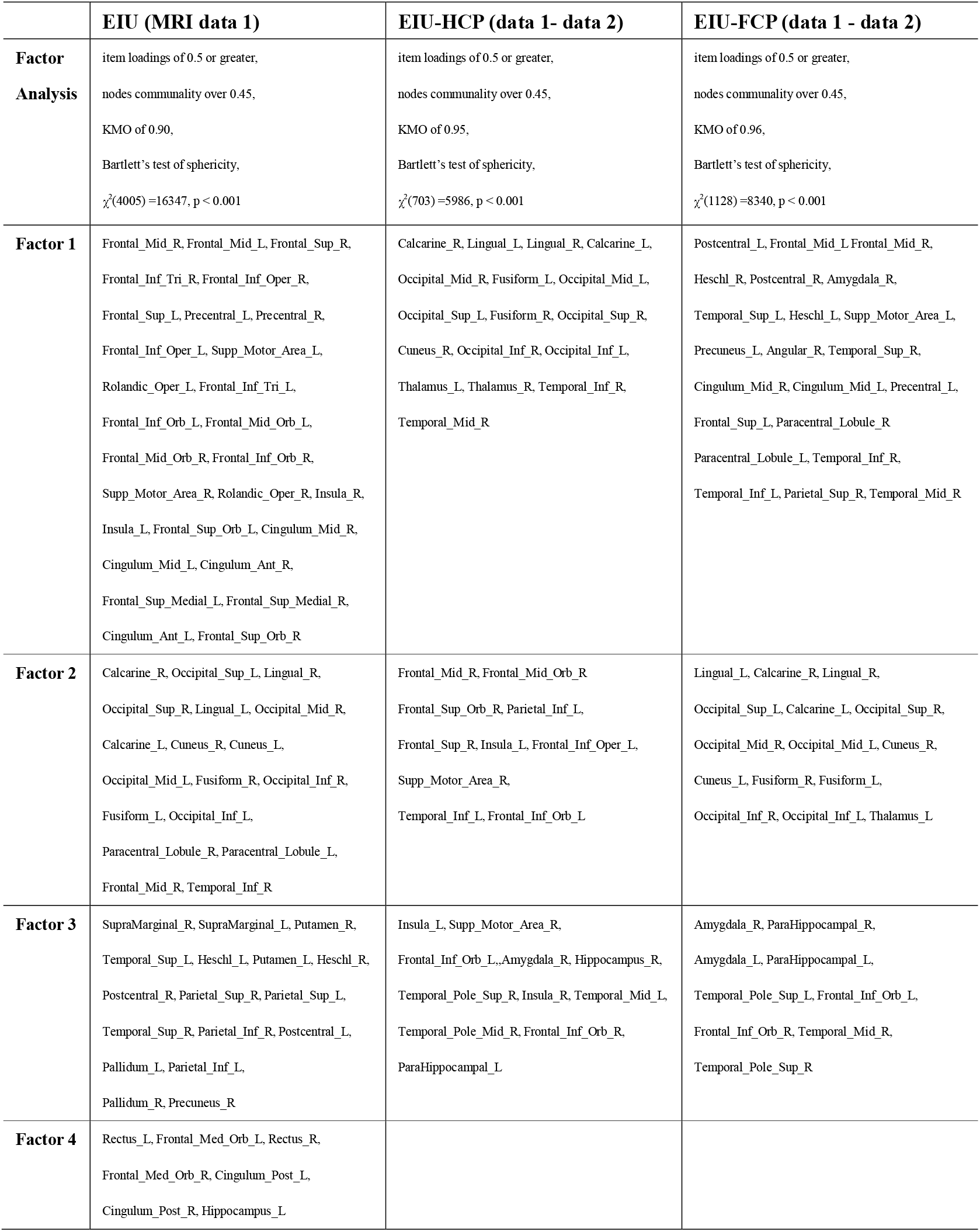
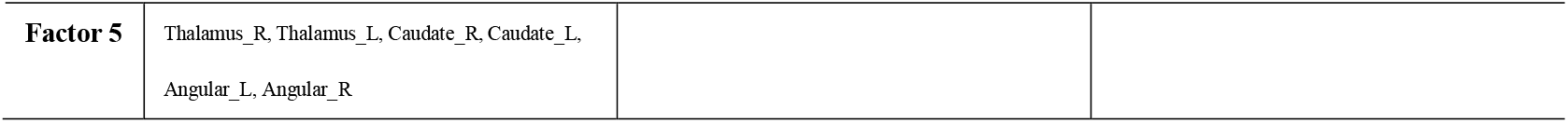
Factor analyses with nodes in three MRI samples.

The factor analysis with the 48 nodes identified 3 major neural networks: factor 1. the frontal-temporal; factor 2. the occipital; and factor 3. the temporal-amygdala. The WBSI total score was negatively correlated with the factor score of the occipital network (Pearson’s r = −0.24, p < 0.05). There was no significant correlation between BIS scores with these networks, ps > 0.05 (Table 3).

Combining the results above, the WBSI scores were positively correlated to the factor scores of the occipital networks in all three analyses. We then extracted the brain regions involved in all three occipital networks, they were the occipital lobe (left and right), lingual lobe (left and right), calcarine (left and right), cuneus (left and right), fusiform (left and right), and left thalamus (Figure 2).

### 4. Identifying the Associated Brain Morphometric Changes

To further clarify the relationship between thought suppression and the occipital lobe, we obtained the following brain morphometric indexes: number of vertices, surface area, gray matter volume, average thickness, integrated rectified mean curvature, integrated rectified gaussian curvature, folding index, and intrinsic curvature index. Pearson’s correlation analyses (p < 0.05) found that the morphometric changes of the medial orbital frontal, supramaginal, banks of superior temporal sulcus, lateral orbito-frontal, superior frontal, rostral middle frontal and transverse temporal lobe were associated with impulsivity (Table 4). The morphometric changes of the caudal anterior cingulum, medial orbital frontal, cuneus, medial orbital frontal, superior frontal, superior parietal, paracentral, precentral, caudal middle frontal, banks of superior temporal sulcus, inferior temporal, temporal pole, middle temporal and entorhinal lobe were associated with thought suppression (Table 4). Overall, the intrinsic curvature index of the left medial orbito-frontal lobe, the number of vertices of the right banks of superior temporal sulcus and the average thickness of the right superior frontal lobe were correlated with thought suppression and impulsivity simultaneously, with same directions. For correlation directions, see Table 4.

**Table 4:** Thought suppression and impulsivity associated brain morphometric changes.

| Index | BIS | WBSI | Overlapped |
| --- | --- | --- | --- |
|  | Brain Region (Corr. R) | Brain Region (Corr. R) | Brain Region |
| **Left:** |  |  |  |
| Intrinsic curvature index | Medial orbito-frontal(-.20) | Caudal Anterior Cingulum(.21);<br>Medial orbito-frontal<br>(-.25) | Medial orbito-frontal |
| Folding index | n/a | n/a | n/a |
| Gray matter volume | n/a | Cuneus(.21) | n/a |
| Integrated rectified<br>gaussian curvature | Supramaginal(-.24) | Caudal Anterior Cingulum(.18) | n/a |
| Number of vertices | n/a | n/a | n/a |
| Integrated rectified<br>mean curvature | Supramaginal(-.30) | Medial orbito-frontal(-.20); Superior<br>parietal(-.20) | n/a |
| Surface area | n/a | Paracentral(-.18); Precentral(-.20) | n/a |
| Average thickness | n/a | Caudal middle frontal(.20) | n/a |
| **Right:** |  |  |  |
| Intrinsic curvature index | Banks of superior temporal<br>sulcus(-.20); | n/a | n/a |
| Folding index | Banks of superior temporal<br>sulcus(-.19); Lateral<br>orbito-frontal(.19) | n/a | n/a |
| Gray matter volume | n/a | Banks of superior temporal<br>sulcus(-.19); Inferior temporal(.18);<br>Temporal pole(.25) | n/a |
| Integrated rectified<br>gaussian curvature | Lateral orbito-frontal(.18) | n/a | n/a |
| Number of vertices | Banks of superior temporal<br>sulcus(-.20) | Banks of superior temporal<br>sulcus(-.19) | Banks of superior temporal sulcus |
| Integrated rectified<br>mean curvature | Banks of superior temporal<br>sulcus(-.22); | n/a | n/a |
| Surface area | n/a | Banks of superior temporal<br>sulcus(-.18); Temporal pole(.18) | n/a |
| Average thickness | Superior frontal(.23);<br>Rostral middle frontal(.21) ;<br>Transverse temporal(-.24) | Superior frontal(.25);<br>Entorhinal(-.23); Middle<br>temporal(.19); Precentral(.18) | Superior frontal |

## DISCUSSION

### 1. Impulsivity and Thought Suppression Share Common Connections

The dysfunction of the top-down mechanism in the brain was related to multiple psychiatric diseases. Through the whole brain analysis of the resting-state functional connectivity and the repetition with different brain atlas, we found that 16 node-to-node connections were associated with thought suppression and impulsivity simultanously, they were divided into 4 groups: 1. the frontal network (the frontal lobe project to the cingulum, hippocampus, parahippocampal, putamen, and angular); 2. the temporal network (the temporal lobe project to the olfactory lobe, hippocampus, parahippocampal gyrus, amygdala, occipital lobe, and parietal lobe); 3. the hippocampus network (the hippocampus project to the middle orbital frontal lobe, inferior orbital frontal lobe, temporal lobe, and amygdala); 4. the cingulum network (the cingulum project to the frontal lobe, parietal lobe, and angular). In addition, the brain morphometric changes related to thought suppression and impulsivity were recorded. The intrinsic curvature index of the left medial orbito-frontal lobe (lower level), the number of vertices of the right banks of superior temporal sulcus (lower level) and the average thickness of the right superior frontal lobe (higher level) were correlated with the higher level of thought suppression and impulsivity. There were extensive functional connections between the frontal lobe and the cingulum, hippocampus, parahippocampal, putamen and angular, and these connections act on both thought suppression and impulsivity. The stronger the connections between the frontal lobe and the related brain regions, the higher levels of thought suppression and impulsivity were. In addition, the frontal lobe, including the left medial orbito-frontal lobe and the right superior frontal lobe, showed the same direction of the change for impulsivity and thought suppression.

### 2. The Occipital Network and Thought Suppression

The study found that the abnormalities in the occipital networkwere strongly associated with the level of thought suppression, but not associated with the morphometric change in the occipital lobe. To our knowledge, this is the first study that revealed the association between the occipital network and thought suppression. The lower level of connection strength in the occipital network was associated with a greater level of thought suppression. With additional rsfMRI data, we found a lower node strength level in the occipital network in excessive internet users compared to healthy participants.

Abnormalities in the occipital lobe have been reported in neuropsychological disorders that were associated with thought suppression. For example, local magnetic low-frequency activity was abnormal in the occipital lobe of untreated depression patients. Increased occipital delta dipole density in depression was a reliable risk factor and was related to disease severity ^57^. The γ-aminobutyric acid (GABA) level in the occipital lobe and ACC of recovered depressed patients were significantly lower than normal controls ^58^. The homogeneity of bilateral occipital lobe was significantly reduced in the resting state of depression, abnormal spontaneous activities in these regions were found in depressed patients ^59^. In patients with major depression, occipital curvature was more common than healthy controls ^60^. The occipital lobes were mainly related to neurocognitive impairment of OCD ^61^. The functional connectivity between the right thalamus and the right middle occipital lobe was related to the patient’s obsessive-compulsive forcing and total score ^62^. The brain regions that exhibited the greatest discrimination in OCD classification included the prefrontal cortex, ACC, anterior central gyrus, and occipital lobe ^63^. Patients with social anxiety disorder had functional connection abnormalities between the frontal and occipital cortex ^64^. State anxiety was positively correlated with functional connectivity between the left lower occipital gyrus and the right medial superior frontal gyrus, and negatively correlated with functional connectivity between the left lower occipital gyrus and the left superior parietal gyrus^65^.

Therefore, the current research was complementary to the past work, by showing the occipital network abnormalities in excessive internet users and its relationship with thought suppression, it further revealed the importance of the occipital network in diseases related to thought suppression.\

### 3. Summary

In the study, the excessive internet users had higher levels of impulsivity and thought suppression than normal population. The two psychological traits were positively correlated with each other and both were related to the psychological symptoms. The study found that thought suppression and impulsivity shared common connections in the top-down mechanism. These common connections maybe a key to study the impulse-inhibition control in the future work.

The study suggested an important role of thought suppression in behavioral addiction, and the relationship between thought suppression and the occipital network, further supported the role of the occipital lobe in psychiatric diseases, the study of the occipital lobe will improve the understanding of psychiatric diseases that related to thought suppression. For the future work, it is important to study not only the control of impulsivity but also the expression of thought suppression in mental illness. This is based on the hypothesis of that the disease may not occur when the two reach to a balance.

## Supporting information

Supplemental for TS full text

## ACKNOWLEDGEMENTS

This work was supported by grants from the National Key Basic Research Program (2016YFA0400900 and 2018YFC0831101), the major Project of Philosophy and Social Science Research, Ministry of Education of China (19JZD010), the National Natural Science Foundation of China (71942003, 31771221, 61773360, and 71874170), the Fundamental Research Funds for the Central Universities of China. A portion of the numerical calculations in this study were performed with the supercomputing system at the Supercomputing Centre of University of Science and Technology of China.

## CONFLICT OF INTEREST

All authors contributed to interpretation of the data, drafting of the manuscript, approval of the final manuscript, and were responsible for the decision to submit the manuscript. All authors report no biomedical financial interests or potential conflicts of interest.

Supplementary information is available at MP’s website.

## REFERENCES

1. Dong G, Potenza MN. A cognitive-behavioral model of Internet gaming disorder: Theoretical underpinnings and clinical implications. Journal of Psychiatric Research 2014; 58:7–11.

2. Goldstein RZ, Volkow ND. Dysfunction of the prefrontal cortex in addiction: neuroimaging findings and clinical implications. Nature Reviews Neuroscience 2011; 12(11):652–669.

3. Brand M, Young KS, Laier C, Woelfling K, Potenza MN. Integrating psychological and neurobiological considerations regarding the development and maintenance of specific Internet-use disorders: An Interaction of Person-Affect-Cognition-Execution (I-PACE) model. Neuroscience and Biobehavioral Reviews 2016; 71:252–266.

4. Wölfling K, Müller KW, Dreier M, Ruckes C, Deuster O, Batra A et al. Efficacy of Short-term Treatment of Internet and Computer Game Addiction: A Randomized Clinical Trial. JAMA Psychiatry 2019; 76(10):1018–1025.

5. Wenzlaff EM, Wegner DM. Thought suppression. Annual Review of Psychology 2000; 51:59–91.

6. Moeller FG, Barratt ES, Dougherty DM, Schmitz JM, Swann AC. Psychiatric aspects of impulsivity. American Journal of Psychiatry 2001; 158(11):1783–1793.

7. Durana M, Gallay R, Robert P, Pruvot FC. NOVEL TYPE SUBMICROMETER RESOLUTION PSEUDORANDOM POSITION OPTICAL ENCODER. Electronics Letters 1993; 29(20):1792–1794.

8. Robbins TW, Gillan CM, Smith DG, de Wit S, Ersche KD. Neurocognitive endophenotypes of impulsivity and compulsivity: towards dimensional psychiatry. Trends in Cognitive Sciences 2012; 16(1):81–91.

9. Grassi G, Pallanti S, Righi L, Figee M, Mantione M, Denys D et al. Think twice: Impulsivity and decision making in obsessive-compulsive disorder. Journal of Behavioral Addictions 2015; 4(4):263–272.

10. Sherman DK, Iacono WG, McGue MK. Attention-deficit hyperactivity disorder dimensions: a twin study of inattention and impulsivity-hyperactivity. Journal of the American Academy of Child and Adolescent Psychiatry 1997; 36(6):745–753.

11. Gut-Fayand A, Dervaux A, Olie JP, Loo H, Poirier MF, Krebs MO. Substance abuse and suicidality in schizophrenia: a common risk factor linked to impulsivity. Psychiatry Research 2001; 102(1):65–72.

12. Molander AC, Mar A, Norbury A, Steventon S, Moreno M, Caprioli D et al. High impulsivity predicting vulnerability to cocaine addiction in rats: some relationship with novelty preference but not novelty reactivity, anxiety or stress. Psychopharmacology 2011; 215(4):721–731.

13. Hoptman MJ, Antonius D, Mauro CJ, Parker EM, Javitt DC. Cortical Thinning, Functional Connectivity, and Mood-Related Impulsivity in Schizophrenia: Relationship to Aggressive Attitudes and Behavior. American Journal of Psychiatry 2014; 171(9):939–948.

14. Ng TH, Stange JP, Black CL, Titone MK, Weiss RB, Abramson LY et al. Impulsivity predicts the onset of DSM-IV-TR or RDC hypomanic and manic episodes in adolescents and young adults with high or moderate reward sensitivity. Journal of Affective Disorders 2016; 198:88–95.

15. Salkovskis PM, Campbell P. Thought suppression induces intrusion in naturally occurring negative intrusive thoughts. Behaviour research and therapy 1994; 32(1):1–8.

16. Palfai TP, Monti PM, Colby SM, Rohsenow DJ. Effects of suppressing the urge to drink on the accessibility of alcohol outcome expectancies. Behaviour research and therapy 1997; 35(1):59–65.

17. Salkovskis PM, Reynolds M. Thought suppression and smoking cessation. Behaviour research and therapy 1994; 32(2):193–201.

18. Gross PR, Eifert GH. Components of generalized anxiety: the role of intrusive thoughts vs worry. Behaviour research and therapy 1990; 28(5):421–428.

19. Kalivas BC, Kalivas PW. Corticostriatal circuitry in regulating diseases characterized by intrusive thinking. Dialogues in Clinical Neuroscience 2016; 18(1):65–76.

20. Collardeau F, Corbyn B, Abramowitz J, Janssen PA, Woody S, Fairbrother N. Maternal unwanted and intrusive thoughts of infant-related harm, obsessive-compulsive disorder and depression in the perinatal period: study protocol. Bmc Psychiatry 2019; 19.

21. Waters FAV, Badcock JC, Michie PT, Maybery MT. Auditory hallucinations in schizophrenia: intrusive thoughts and forgotten memories. Cognitive neuropsychiatry 2006; 11(1):65–83.

22. Bomyea J, Lang AJ. Accounting for intrusive thoughts in PTSD: Contributions of cognitive control and deliberate regulation strategies. Journal of Affective Disorders 2016; 192:184–190.

23. Wegner DM, Zanakos S. Chronic thought suppression. Journal of personality 1994; 62(4):616–640.

24. Keough ME, Timpano KR, Riccardi CJ, Schmidt NB. Suppressing the white bears interacts with anxiety sensitivity in the prediction of mood and anxiety symptoms. Personality and Individual Differences 2010; 49(5):408–413.

25. Magee JC, Zinbarg RE. Suppressing and focusing on a negative memory in social anxiety: Effects on unwanted thoughts and mood. Behaviour Research and Therapy 2007; 45(12):2836–2849.

26. Julien D, O’Connor KP, Aardema F. Intrusive thoughts, obsessions, and appraisals in obsessive-compulsive disorder: A critical review. Clinical Psychology Review 2007; 27(3):366–383.

27. Wenzlaff RM, Bates DE. Unmasking a cognitive vulnerability to depression: how lapses in mental control reveal depressive thinking. Journal of personality and social psychology 1998; 75(6):1559–1571.

28. Haber SN, Fudge JL, McFarland NR. Striatonigrostriatal pathways in primates form an ascending spiral from the shell to the dorsolateral striatum. Journal of Neuroscience 2000; 20(6):2369–2382.

29. Buckner RL, Andrews-Hanna JR, Schacter DL. The brain’s default network - Anatomy, function, and relevance to disease. In: Kingstone A, Miller MB (eds). Year in Cognitive Neuroscience 2008, vol. 11242008, pp 1–38.

30. Bonelli RM, Cummings JL. Frontal-subcortical circuitry and behavior. Dialogues in clinical neuroscience 2007; 9(2):141–151.

31. Dalley JW, Everitt BJ, Robbins TW. Impulsivity, Compulsivity, and Top-Down Cognitive Control. Neuron 2011; 69(4):680–694.

32. Korponay C, Pujara M, Deming P, Philippi C, Decety J, Kosson DS et al. Impulsive-antisocial psychopathic traits linked to increased volume and functional connectivity within prefrontal cortex. Social Cognitive and Affective Neuroscience 2017; 12(7):1169–1178.

33. Beck SR, Riggs KJ, Gorniak SL. Relating developments in children’s counterfactual thinking and executive functions. Thinking & Reasoning 2009; 15(4):337–354.

34. Ziegler G, Hauser TU, Moutoussis M, Bullmore ET, Goodyer IM, Fonagy P et al. Compulsivity and impulsivity traits linked to attenuated developmental frontostriatal myelination trajectories. Nature Neuroscience 2019; 22(6):992–+.

35. Ma L, Steinberg JL, Cunningham KA, Bjork JM, Lane SD, Schmitz JM et al. Altered anterior cingulate cortex to hippocampus effective connectivity in response to drug cues in men with cocaine use disorder. Psychiatry Research-Neuroimaging 2018; 271:59–66.

36. Fujihara K, Narita K, Suzuki Y, Takei Y, Suda M, Tagawa M et al. Relationship of gamma-aminobutyric acid and glutamate plus glutamine concentrations in the perigenual anterior cingulate cortex with performance of Cambridge Gambling Task. Neuroimage 2015; 109:102–108.

37. Passamonti L, Rowe JB, Schwarzbauer C, Ewbank MP, von dem Hagen E, Calder AJ. Personality Predicts the Brain’s Response to Viewing Appetizing Foods: The Neural Basis of a Risk Factor for Overeating. Journal of Neuroscience 2009; 29(1):43–51.

38. Mitchell JP, Heatherton TF, Kelley WM, Wyland CL, Wegner DM, Neil MacRae C. Separating sustained from transient aspects of cognitive control during thought suppression. Psychological Science 2007; 18(4):292–297.

39. Mitchell JP, Heatherton TF, Kelley WM, Wyland CL, Wegner DM, Macrae CN. Separating sustained from transient aspects of cognitive control during thought suppression. Psychological Science 2007; 18(4):292–297.

40. Aso T, Nishimura K, Kiyonaka T, Aoki T, Inagawa M, Matsuhashi M et al. Dynamic interactions of the cortical networks during thought suppression. Brain and Behavior 2016; 6(8).

41. Tsujii N, Mikawa W, Tsujimoto E, Adachi T, Niwa A, Ono H et al. Reduced left precentral regional responses in patients with major depressive disorder and history of suicide attempts. Plos One 2017; 12(4).

42. Xu S, Li M, Yang C, Fang X, Ye M, Wei L et al. Altered Functional Connectivity in Children With Low-Function Autism Spectrum Disorders. Frontiers in Neuroscience 2019; 13.

43. Bullmore ET, Sporns O. Complex brain networks: graph theoretical analysis of structural and functional systems. Nature Reviews Neuroscience 2009; 10(3):186–198.

44. Damoiseaux JS, Rombouts SARB, Barkhof F, Scheltens P, Stam CJ, Smith SM et al. Consistent resting-state networks across healthy subjects. Proceedings of the National Academy of Sciences of the United States of America 2006; 103(37):13848–13853.

45. Hull JV, Jacokes ZJ, Torgerson CM, Irimia A, Van Horn JD, Consortium GR. Resting-State Functional Connectivity in Autism Spectrum Disorders: A Review. Frontiers in Psychiatry 2017; 7,205.

46. Wang L, Hermens DF, Hickie IB, Lagopoulos J. A systematic review of resting-state functional-MRI studies in major depression. Journal of Affective Disorders 2012; 142(1-3):6–12.

47. Sun D-L, Chen Z-J, Ma N, Zhang X-C, Fu X-M, Zhang D-R. Decision-Making and Prepotent Response Inhibition Functions in Excessive Internet Users. Cns Spectrums 2009; 14(2):75–81.

48. Derogatis LR. Symptom Checklist-90-R [SCL-90-R]Administration, Scoring, and Procedures Manual (3^rd^ ed.). National Computer Systems: Minneapolis, MN,1994.

49. Patton JH, Stanford MS, Barratt ES. Factor structure of the Barratt impulsiveness scale. Journal of clinical psychology 1995; 51(6):768–774.

50. Mas-Exposito L, Amador-Campos JA, Gomez-Benito J, Lalucat-Jo L. The World Health Organization Quality of Life Scale Brief Version: a validation study in patients with schizophrenia. Quality of Life Research 2011; 20(7):1079–1089.

51. Cloninger CR, Przybeck TR, Svrakic DM. The Tridimensional Personality Questionnaire: U.S. normative data. Psychological reports 1991; 69(3 Pt 1):1047–1057.

52. Bullmore ET, Bassett DS. Brain Graphs: Graphical Models of the Human Brain Connectome. In: NolenHoeksema S, Cannon TD, Widiger T (eds). Annual Review of Clinical Psychology, vol. 72011, pp 113–140.

53. Tzourio-Mazoyer N, Landeau B, Papathanassiou D, Crivello F, Etard O, Delcroix N et al. Automated anatomical labeling of activations in SPM using a macroscopic anatomical parcellation of the MNI MRI single-subject brain. Neuroimage 2002; 15(1):273–289.

54. Fan L, Li H, Zhuo J, Zhang Y, Wang J, Chen L et al. The Human Brainnetome Atlas: A New Brain Atlas Based on Connectional Architecture. Cerebral Cortex 2016; 26(8):3508–3526.

55. Whelan R, Conrod PJ, Poline JB, Lourdusamy A, Banaschewski T, Barker GJ et al. Adolescent impulsivity phenotypes characterized by distinct brain networks. Nature Neuroscience 2012; 15(6):920–925.

56. Desikan RS, Segonne F, Fischl B, Quinn BT, Dickerson BC, Blacker D et al. An automated labeling system for subdividing the human cerebral cortex on MRI scans into gyral based regions of interest. Neuroimage 2006; 31(3):968–980.

57. Fernandez A, Rodriguez-Palancas A, Lopez-Ibor M, Zuluaga P, Turrero A, Maestu F et al. Increased occipital delta dipole density in major depressive disorder determined by magnetoencephalography. Journal of Psychiatry & Neuroscience 2005; 30(1):17–23.

58. Bhagwagar Z, Wylezinska M, Jezzard P, Evans J, Boorman E, Matthews PM et al. Low GABA concentrations in occipital cortex and anterior cingulate cortex in medication-free, recovered depressed patients. International Journal of Neuropsychopharmacology 2008; 11(2):255–260.

59. Peng D-h, Jiang K-d, Fang Y-r, Xu Y-f, Shen T, Long X-y et al. Decreased regional homogeneity in major depression as revealed by resting-state functional magnetic resonance imaging. Chinese Medical Journal 2011; 124(3):369–373.

60. Maller JJ, Thomson RHS, Rosenfeld JV, Anderson R, Daskalakis ZJ, Fitzgerald PB. Occipital bending in depression. Brain 2014; 137:1830–1837.

61. Wen S-l, Cheng M-f, Cheng M-h, Yue J-h, Li J-f, Xie L-j. Neurocognitive Dysfunction and Regional Cerebral Blood Flow in Medically Na ve Patients With Obsessive-Compulsive Disorder. Developmental Neuropsychology 2014; 39(1):37–50.

62. Chen Y, Meng Z, Zhang Z, Zhu Y, Gao R, Cao X et al. The right thalamic glutamate level correlates with functional connectivity with right dorsal anterior cingulate cortex/middle occipital gyrus in unmedicated obsessive-compulsive disorder: A combined fMRI and H-1-MRS study. Australian and New Zealand Journal of Psychiatry 2019; 53(3):207–218.

63. Bu X, Hu X, Zhang L, Li B, Zhou M, Lu L et al. Investigating the predictive value of different resting-state functional MRI parameters in obsessive-compulsive disorder. Translational Psychiatry 2019; 9:17.

64. Ding J, Chen H, Qiu C, Liao W, Warwick JM, Duan X et al. Disrupted functional connectivity in social anxiety disorder: a resting-state fMRI study. Magnetic Resonance Imaging 2011; 29(5):701–711.

65. Li K, Zhang M, Zhang H, Li X, Zou F, Wang Y et al. The spontaneous activity and functional network of the occipital cortex is correlated with state anxiety in healthy adults. Neuroscience Letters 2020; 715:134596.

