## Supplemental for TS full text for "Impulsivity and Thought Suppression in Behavioral Addiction: Associated Neural Connectivity and Neural Networks"

**Supplementary**

A. **Supplemental Experimental Procedure**
1. **Participants**
2. **Assessment**
3. **Resting-state fMRI Data Acquisition**
4. **Pre-processing and Functional Connectivity**
5. **T1 Structural Image Data Analysis**
B. **Supplemental Tables**

**Table S1: Psychological Measures**

**Table S2: Nine items in the Symptom Checklist 90-R: BIS 2 group x WBSI 2 group**

**Table S3: The correlations between BIS/WBSI scores and neural connections**

C. **Supplemental Reference**
A. **Supplemental Experimental Procedure**
1. **Participants**

Data were acquired from 131 internet users. Participants who meet the following criteria were recruited: 1) internet use for at least 12 months prior to the experiment; 2) continues internet use everyday in the past 3 months, beyond the purpose of work or study; 3) right-handedness; 4) age 14-25 years old; 5) male; 6) no history of or current neurological or psychiatric disorders, including substance use; 7) no medicine use or substance abuse in the past 3 months. The experiment conformed to the Code of Ethics of the World Medical Association (Declaration of Helsinki). The study was approved by the Human Research Ethics Committee of the University of Science and Technology of China. All participants provided written informed consent.

2. **Assessment**
1) Excessive Internet Use ^1^: Excessive internet use referred compulsive online purchasing, gambling, chatting, video streaming, excessive shopping, or aimless information searching ^2^. The questionnaire assess the use of internet, including time, frequency, type and style, in the past one week and the past one year separately. In addition, internet addiction related symptoms were listed, the score is calculated based on the number of symptoms happen in the past three months. Based on empirical evidence, excessive online activities was likely associated with marked functional impairment, including compulsive online buying, gambling, cybersex, as well as excessive use of online streaming and social media that have addictive, impulsive and/or compulsive elements.
2) Symptom Checklist 90-Revised ^3^: The checklist assess whether a person has certain psychological symptoms and how severe it is. All 90 items are divided into 10 dimensions: somatization, obsessive-compulsive, interpersonal sensitivity, depression, anxiety, hostility, phobic anxiety, paranoid ideation, psychoticism, and others. For each dimension, a score of 1.5-2.5 refers individual feel the symptoms, the severity is mild to moderate; a score of 2.5-3.5 refers that individual feel the symptoms, the severity is moderate to severe.
3) Barratt Impulsiveness Scale (BIS; ^4^: To assess the personality/behavioral construct of impulsiveness. The scale produces 6 first-order factors: attention, motor, self-control, cognitive complexity, perseverance and cognitive instability, and 3 second-order factors: attentional , motor, and nonplanning, with the higher score indicating a greater level of impulsiveness. The Chinese version of BIS has been tested, Cronbach’s alpha = 0.67-0.89, and re-test reliability (1-month), r = 0.70, p < 0.001 ^5^.
4) White Bear Suppression Inventory (WBSI; ^6^: WBSI is a 15-item questionnaire that measures thought suppression. There are three dimensions: intrusive thoughts, thought suppression, and attention switch. The Chinese version of the WBSI has been confirmed to have good reliability and validity, Cronbach’s alpha = 0.87, and re-test reliability (1-month), r = 0.59, p < 0.001 ^7^.
5) World Health Organization Quality of Life Scale-Brief Form Questionnaire (QoL; WHO, 1991): It assesses the individual's perceptions in the context of their culture and value systems, and their personal goals, standards and concerns. The aim was to develop an international cross-culturally comparable quality of life assessment instrument. The instrument was developed collaboratively in a number of centers worldwide, and have been widely field-tested. The instrument comprises 26 items, which measure the following broad domains: physical health, psychological health, social relationships, and environment.
6) Tridimensional Personality Questionnaire (TPQ) ^8^: TPQ is a 60-item questionnaire that measures personality traits.The questionnaire has twelve factors in three dimensions: novelty seeking (seek excitement and rigidity, impulsive, unscrupulous, conservative in words and deeds, unruly, follow the rules); harm avoidance (anticipated anxiety and pessimism, unconstrained optimism, fear of uncertainty, afraid of meeting strangers/shy, fatigue/weakness; reward dependence (sentimentality, perseverance, attachment, dependence).
3. **Resting-state fMRI Data Acquisition**

MRI data were acquired using two identical 3-T Siemens Magnetom Trio scanners (Siemens, Erlangen, Germany) on two sites (Anhui Provincial Hospital and General Hospital of Beijing Military Region). Effect of scan site was controlled by adding it as a nuisance co-variate in all statistical analyses. A circularly polarized head coil was used, with foam padding to restrict head motion. Functional images were acquired with a T2-weighted gradient echo-planar imaging sequence (TR=2 s, TE=30 ms, FOV=240 mm, matrix, 64×64) with 33 axial slices (3.7-mm thickness, no gap), covering the whole brain. All participants were told to keep their heads steady during all runs before entering the MR scanner. High resolution T1-weighted spin-echo images were also collected for anatomical overlay. Resting-state fMRI data were acquired with a 8-min run (240 epochs) while participants were instructed to remain still, keep their eyes closed, not to think of anything and not to fall into sleep.

Additional fMRI data were downloaded from two sources: the Human Connectome Project (HCP; http://humanconnectome.org), and the 1000 Functional Connectomes Project (FCP, [http://fcon_1000.projects.nitrc.org).](https://nunda.northwestern.edu).) The HCP dataset contained 134 individuals (age range: 18-26 years old; males) and the FCP dataset contained 75 individuals (age range: 18-26 years old; males). Both datasets were acquired using gradient-echo echo-planar-imaging on 3 Tesla scanners. For HCP dataset, brain volumes were acquired using the following parameters: TR = 720ms, voxel size = 2×2×2mm^3^ , FOV = 208×180mm^2^, 72 slices, TE = 33.1ms, flip angle = 52 degrees; For the FCP dataset, brain volumes were acquired using the following parameters: TR = 2000ms, voxel size = 3.125×3.125×3.600mm^3^, 33 slices.

4. **Pre-processing and Functional Connectivity**

The imaging data were processed with the Analysis of Functional Neuroimages (Version AFNI_18.2.03). The first two volumes of each run were discarded. Volumes were corrected for temporal shifts between slices, corrected for motion. Using linear regression, we performed linear trend removal. Then, the fMRI data were transformed non-linearly to Montreal Neurological Institute (MNI) 152 space (resampled voxel size 3.5x3.5x3.5mm^3^) according to the spatial transformation between the anatomical data and the MNI space. Finally, volumes were spatially smoothed with an 8-mm full-width at half-maximum Gaussian kernel and grand-mean scaled.

We have used the automated anatomical labeling atlas (AAL) to divide the brain into 90 brain regions to calculate functional connectivity. First, correlations were calculated based on the average time course in each AAL parcel (as a node) as functional connections between brain regions. Thus, each subject generate a 90x90 functional connectivity matrix, which was next used to calculate node strength. Node strength is a measure that quantifies node importance/centrality through the connectivity strength between nodes and all other nodes in a network ^9^.We quantify the strength of a node by calculating the sum of weights of links connected to the node.

5. **T1 Structural Image Data Analysis**

T1 structural image data were analyzed using FreeSurfer (version 6.0), with Desikan-Killiany Atlas. In the cortical surface stream, the tools constructed models of the boundary between white matter and cortical gray matter as well as the pial surface. Cortical surface reconstruction was performed with standard procedures provided by the software (http://surfer.nmr.mgh.harvard.edu/fswiki). Then we obtained an array of anatomical measures, including: cortical thickness, surface area, curvature, and surface normal at each point on the cortex.

B. **Supplemental Tables**

**Table S1: Psychological Measures (n =131)**

| **Items** | **Min** | **Max** | **Mean** | **SD** |
| --- | --- | --- | --- | --- |
| Intrusive thinking | 10.00 | 35.00 | 24.71 | 5.46 |
| Suppressed thinking | 6.00 | 20.00 | 13.29 | 2.77 |
| Attention switch | 3.00 | 15.00 | 9.84 | 2.33 |
| WBSI total score | 22.00 | 70.00 | 47.84 | 9.30 |
| EIU score1 in the worst week | 0.00 | 37.00 | 10.44 | 10.29 |
| EIU score2 in the worst week | 0.00 | 9.00 | 1.80 | 2.46 |
| EIU score1 in the past year | 0.00 | 63.00 | 20.26 | 13.98 |
| EIU score2 in the past year | 0.00 | 9.00 | 4.42 | 3.48 |
| Somatization | 1.00 | 2.80 | 1.44 | 0.44 |
| Obsessive-compulsive symptoms | 1.00 | 3.44 | 1.75 | 0.51 |
| Interpersonal sensitivity | 1.00 | 3.22 | 1.66 | 0.48 |
| Depression | 1.00 | 2.92 | 1.54 | 0.43 |
| Anxiety | 1.00 | 3.00 | 1.38 | 0.37 |
| Hostility | 1.00 | 3.50 | 1.48 | 0.52 |
| Terror | 1.00 | 2.50 | 1.27 | 0.30 |
| Paranoid | 1.00 | 2.83 | 1.43 | 0.43 |
| Psychosis | 1.00 | 2.40 | 1.38 | 0.33 |
| SCL-90 averaged score | 0.99 | 2.46 | 1.47 | 0.32 |
| Physiology | 66.67 | 100 | 84.29 | 9.25 |
| Psychology | 29.41 | 100 | 71.94 | 13.89 |
| Environment | 33.33 | 100 | 65.56 | 12.53 |
| WHO-life total score | 20.00 | 100 | 69.89 | 14.90 |
| TPQ hunting | 3.00 | 28.00 | 14.44 | 5.13 |
| TPQ dodging | 4.00 | 31.00 | 14.18 | 6.06 |
| TPQ reward dependence | 11.00 | 29.00 | 18.35 | 3.66 |
| Attention | 11.00 | 26.00 | 16.72 | 2.89 |
| Motivation | 15.00 | 32.00 | 21.40 | 3.32 |
| Unplanned | 16.00 | 36.00 | 25.26 | 4.47 |
| BIS total score | 47.00 | 85.00 | 63.38 | 8.47 |

**Table S2: Nine items in the Symptom Checklist 90-R: BIS 2 group x WBSI 2 group (0.00 is low score group, 1.00 is high score group)**

| **Symptoms** | **BIS**  **group** | **WBSI group** | **Mean** | **SD** | **N** |
| --- | --- | --- | --- | --- | --- |
| Somatization | 0.00 | 0.00 | 1.27 | 0.32 | 34.00 |
|  |  | 1.00 | 1.48 | 0.48 | 23.00 |
|  | 1.00 | 0.00 | 1.40 | 0.33 | 26.00 |
|  |  | 1.00 | 1.60 | 0.51 | 37.00 |
| Obsessive-compulsive | 0.00 | 0.00 | 1.45 | 0.37 |  |
|  |  | 1.00 | 1.72 | 0.56 |  |
|  | 1.00 | 0.00 | 1.73 | 0.35 |  |
|  |  | 1.00 | 2.05 | 0.54 |  |
| Interpersonal sensitivity | 0.00 | 0.00 | 1.36 | 0.28 |  |
|  |  | 1.00 | 1.67 | 0.54 |  |
|  | 1.00 | 0.00 | 1.59 | 0.37 |  |
|  |  | 1.00 | 1.95 | 0.48 |  |
| Depression | 0.00 | 0.00 | 1.28 | 0.24 |  |
|  |  | 1.00 | 1.44 | 0.32 |  |
|  | 1.00 | 0.00 | 1.62 | 0.38 |  |
|  |  | 1.00 | 1.77 | 0.50 |  |
| Anxiety | 0.00 | 0.00 | 1.20 | 0.22 |  |
|  |  | 1.00 | 1.41 | 0.36 |  |
|  | 1.00 | 0.00 | 1.36 | 0.26 |  |
|  |  | 1.00 | 1.53 | 0.48 |  |
| Hostility | 0.00 | 0.00 | 1.26 | 0.46 |  |
|  |  | 1.00 | 1.57 | 0.55 |  |
|  | 1.00 | 0.00 | 1.44 | 0.37 |  |
|  |  | 1.00 | 1.66 | 0.58 |  |
| Terror | 0.00 | 0.00 | 1.13 | 0.18 |  |
|  |  | 1.00 | 1.28 | 0.32 |  |
|  | 1.00 | 0.00 | 1.26 | 0.24 |  |
|  |  | 1.00 | 1.41 | 0.36 |  |
| Paranoid | 0.00 | 0.00 | 1.20 | 0.26 |  |
|  |  | 1.00 | 1.48 | 0.46 |  |
|  | 1.00 | 0.00 | 1.31 | 0.36 |  |
|  |  | 1.00 | 1.68 | 0.46 |  |
| Psychosis | 0.00 | 0.00 | 1.18 | 0.19 |  |
|  |  | 1.00 | 1.45 | 0.33 |  |
|  | 1.00 | 0.00 | 1.32 | 0.28 |  |
|  |  | 1.00 | 1.53 | 0.36 |  |

**Table S3: The correlations between BIS score (WBSI score) and neural connection, by using two brain templates (AAL and Brainnetome).**

| **AAL template** | **BIS** | **WBSI** | **BIS** | **WBSI** | **Brainnetome template** | **BIS** | **WBSI** | **BIS** | **WBSI** |
| --- | --- | --- | --- | --- | --- | --- | --- | --- | --- |
| **Connections** | **r*** | **r** | **p** | **p** | **Connections** | **r** | **r** | **p** | **p** |
| Frontal_Mid_R-Cingulum_Post_L | 0.18 | 0.22 | 0.05 | 0.02 | MFG_R_7_3-CG_L_7_1 | 0.18 | 0.22 | 0.05 | 0.02 |
| Frontal_Mid_R-Cingulum_Post_R | 0.19 | 0.23 | 0.04 | 0.01 | MFG_R_7_3-CG_R_7_1 | 0.21 | 0.21 | 0.02 | 0.02 |
| Frontal_Mid_Orb_R-Hippocampus_L | 0.20 | 0.20 | 0.03 | 0.03 | OrG_R_6_6-Hipp_L_2_1 | 0.20 | 0.17 | 0.03 | 0.05 |
| Frontal_Mid_Orb_R-Hippocampus_R | 0.18 | 0.19 | 0.05 | 0.03 | OrG_R_6_6-Hipp_R_2_2 | 0.20 | 0.15 | 0.03 | 0.05 |
| Frontal_Inf_Tri_R-Cingulum_Post_R | 0.18 | 0.20 | 0.04 | 0.03 | IFG_R_6_4-CG_R_7_4 | 0.20 | 0.24 | 0.03 | 0.01 |
| Frontal_Inf_Orb_L-ParaHippocampal_L | 0.20 | 0.27 | 0.03 | 0.00 | OrG_L_6_6-PhG_L_6_3 | 0.19 | 0.17 | 0.04 | 0.05 |
| Frontal_Inf_Orb_R-ParaHippocampal_R | 0.23 | 0.18 | 0.01 | 0.05 | SFG_R_7_7-Str_L_6_4 | 0.23 | 0.19 | 0.01 | 0.04 |
| Olfactory_L-Temporal_Mid_R | 0.21 | 0.19 | 0.02 | 0.03 | SPL_L_5_3-CG_L_7_1 | 0.22 | 0.17 | 0.02 | 0.05 |
| Frontal_Sup_Medial_L-Putamen_L | 0.21 | 0.19 | 0.02 | 0.04 | IPL_R_6_4-CG_L_7_4 | 0.17 | 0.18 | 0.05 | 0.05 |
| Frontal_Sup_Medial_R-Putamen_L | 0.18 | 0.22 | 0.05 | 0.02 | STG_L_6_3-Hipp_R_2_1 | 0.18 | 0.18 | 0.05 | 0.04 |
| Cingulum_Post_L-Parietal_Sup_L | 0.19 | 0.23 | 0.04 | 0.01 | PhG_L_6_5-Amyg_L_2_1 | 0.17 | 0.17 | 0.05 | 0.05 |
| Cingulum_Post_L-Angular_R | 0.20 | 0.19 | 0.03 | 0.04 | STG_L_6_3-PhG_L_6_6 | 0.18 | 0.17 | 0.05 | 0.04 |
| Hippocampus_L-Temporal_Sup_L | 0.18 | 0.21 | 0.04 | 0.02 | STG_L_6_4-Amyg_L_2_1 | 0.18 | 0.18 | 0.04 | 0.04 |
| Hippocampus_R-Temporal_Sup_L | 0.19 | 0.23 | 0.04 | 0.01 | STG_L_6_6-SPL_L_5_3 | 0.18 | 0.18 | 0.05 | 0.04 |
| ParaHippocampal_L-Amygdala_L | 0.20 | 0.24 | 0.03 | 0.01 | STG_L_6_4-ITG_L_7_5 | 0.23 | 0.18 | 0.01 | 0.05 |
| ParaHippocampal_L-Temporal_Pole_Sup_L | 0.20 | 0.26 | 0.03 | 0.00 |  |  |  |  |  |
| Amygdala_L-Temporal_Pole_Sup_L | 0.22 | 0.18 | 0.02 | 0.04 |  |  |  |  |  |
| Occipital_Inf_L-Temporal_Mid_L | 0.19 | 0.21 | 0.04 | 0.02 |  |  |  |  |  |
| Parietal_Sup_L-Temporal_Pole_Sup_L | 0.20 | 0.19 | 0.03 | 0.04 |  |  |  |  |  |
| Temporal_Pole_Sup_L-Temporal_Inf_L | 0.21 | 0.20 | 0.02 | 0.03 |  |  |  |  |  |

*r is correlation coefficient

Abbreviation: MFG, Middle Frontal Gyrus; CG, Cingulate Gyrus; OrG, Orbital Gyrus; PhG, Parahippocampal Gyrus; STG, Superior Temporal Gyrus; SPL, Superior Parietal Lobule; IPL, Inferior Parietal Lobule; TG, Inferior Temporal Gyrus; ITG, Inferior Temporal Gyrus; Str, striatum; Hipp, hippocampus; Amyg, amygdala.

C. **References**
1. Cloninger CR, Przybeck TR, Svrakic DM. The Tridimensional Personality Questionnaire: U.S. normative data. *Psychological Reports* 1991; 69(3 Pt 1): 1047-57.
2. Dale AM, Fischl B, Sereno MI. Cortical surface-based analysis - I. Segmentation and surface reconstruction. *Neuroimage* 1999; 9(2): 179-94.
3. Derogatis LR. Symptom Checklist-90-R [SCL-90-R]：Administration, Scoring, and Procedures Manual (3rdedn). National Computer Systems: Minneapolis, MN,1994
4. Desikan RS, Segonne F, Fischl B, Quinn BT, Dickerson BC, Blacker D, et al. An automated labeling system for subdividing the human cerebral cortex on MRI scans into gyral based regions of interest. *Neuroimage* 2006; 31(3): 968-80.
5. Fischl B, Sereno MI, Dale AM. Cortical surface-based analysis - II: Inflation, flattening, and a surface-based coordinate system. *Neuroimage* 1999; 9(2): 195-207.
6. Wegner DM, Zanakos S. Chronic Thought Suppression. *Journal of Personality* 1994; 62(4): 615-640.
7. Reyes-Rodríguez ML, Rivera-Medina CL, Cámara-Fuentes L, Suárez-Torres A, Bernal G. Depression symptoms and stressful life events among college students in Puerto Rico. *Journal of Affective Disorders* 2013; 145(3): 324-330.
8. Cloninger CR, Przybeck TR, Svrakic DM. The Tridimensional Personality Questionnaire: U.S. normative data. *Psychological Reports* 1991; 69(3 Pt 1): 1047-1057.
9. Rubinov M, Sporns O. Complex network measures of brain connectivity: Uses and interpretations. *Neuroimage* 2010; 52(3): 1059-1069.
10. Zou Q-H, Zhu C-Z, Yang Y, Zuo X-N, Long X-Y, Cao Q-J et al. An improved approach to detection of amplitude of low-frequency fluctuation (ALFF) for resting-state fMRI: Fractional ALFF. *Journal of Neuroscience Methods* 2008; 172(1): 137-141.
11. Desikan RS, Segonne F, Fischl B, Quinn BT, Dickerson BC, Blacker D et al. An automated labeling system for subdividing the human cerebral cortex on MRI scans into gyral based regions of interest. *Neuroimage* 2006; 31(3): 968-980.
12. A. M. Dale BF, and M. I. Sereno. Cortical surface-based analysis i. segmentation and surface reconstruction. *Neuroimage* 1999; 9(2): 179-194.
13. B. Fischl MIS, and A. M. Dale. Cortical surface-based analysis ii: Inflation, flattening, and a surfacebased coordinate system. *Neuroimage* 1999; 9(2): 195-207.
